# A Mean Shift Algorithm for Drift Correction in Localization Microscopy

**DOI:** 10.1101/2021.05.07.443176

**Authors:** Frank J Fazekas, Thomas R Shaw, Sumin Kim, Ryan A Bogucki, Sarah L Veatch

## Abstract

Single molecule localization microscopy (SMLM) techniques transcend the diffraction limit of visible light by localizing isolated emitters sampled stochastically. This time-lapse imaging necessitates long acquisition times, over which sample drift can become large relative to the localization precision. Here we present a novel, efficient, and robust method for estimating drift using a simple peak-finding algorithm based on mean shifts that is effective for SMLM in 2 or 3 dimensions.

## Main

Stochastic super-resolution microscopy techniques such as STORM ^1,2^ and PALM ^3,4^ exploit photoswitching of fluorescent probes to enable imaging of densely labeled samples with resolutions an order of magnitude smaller than the diffraction limit of visible light. Sparsely distributed point spread functions (PSFs) of single emitters are identified in individual image frames, and their centroids are determined according to an appropriate fitting algorithm. The final reconstruction is typically a 2D or 3D histogram of these single-molecule positions. Drift due to thermal expansion or mechanical instabilities can degrade image quality over the course of image acquisition, which typically occurs on the timescale of minutes. Drift compensation requires either active stabilization of the microscope or *a posteriori* computation of the drift curves either using fiducial markers or the acquired single molecule localizations. In this report, we present a mathematically simple approach to drift correction using a mean shift (MS) algorithm ^5–7^ for static SMLM datasets without fiducial markers, with some advantages over past approaches that use nonlinear least squares (NLLS) fitting of image-based cross-correlations ^8–10^.

A graphical illustration of the mean shift algorithm as applied to sample 2D localizations is presented in Fig 1. The localizations all lie in one of two datasets which sample the same structure with a constant relative shift **r**_shift_ in space. The first step of the algorithm is to extract pairwise displacements between all localizations across the two datasets. When individual displacements are plotted as points (Fig 1b), displacements arising from the same labeled objects (magenta points) cluster around **r**_shift_, while displacements arising from different objects (green points) distribute randomly over space. The mean shift algorithm determines the center of the peak of the distribution of through iteration ^5–7^. At each iteration, all pairs within the radius of consideration are extracted, and the updated shift estimate is the centroid of these pairs. The uniformly distributed background will tend to bias the centroid towards the center of the observation window, while the peak moves the mean toward **r**_shift_. The observation window is then redrawn around the new mean and the process is repeated until the peak is centered in the observation window. Three iterations of the algorithm are visualized in Fig 1c.

**Figure 1.**
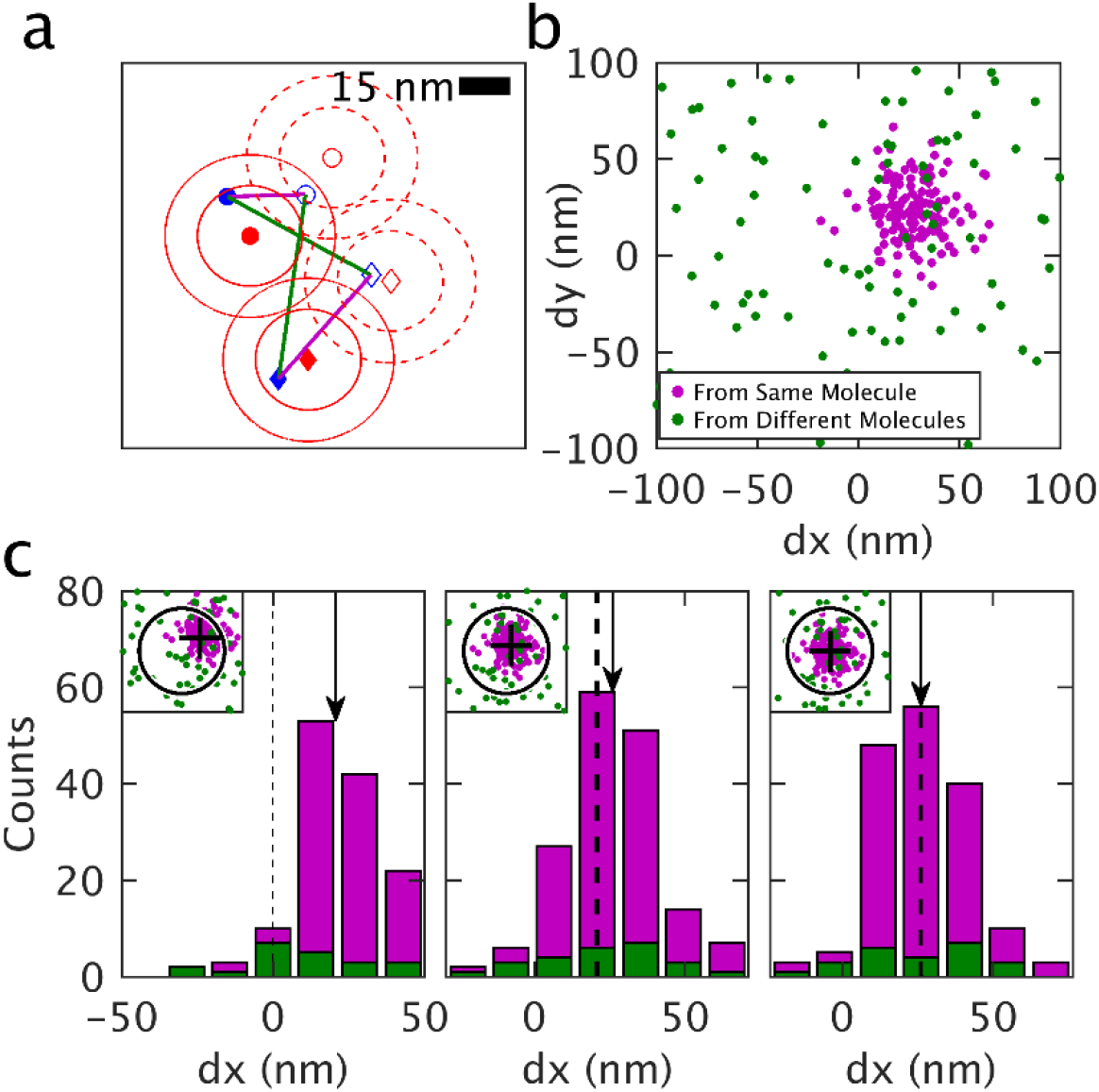
Demonstration of the mean shift algorithm. a) A sample 2D image containing two molecules (filled red symbols) that are each localized one time (filled blue symbols). At a later time the molecules are translated 25nm in both dimensions (open red symbols) and two additional localizations are acquired (open blue symbols). Red contours indicate 1 and 2 times the localization precision around molecule centers (15nm). Straight lines show displacements between localizations acquired at distinct times. Some connect localizations from the same displaced molecule (magenta) while others connect different molecules (green). b) Displacements like those shown in (a) displayed as points. Points connecting the same molecules cluster around the displacement while points connecting different molecules produce a uniform background. c) Three iterations of the mean shift algorithm showing the displacements as histograms in one dimension and as points in 2D in the inset. Initially, a region of interest (circle in inset) is centered at zero shift. The mean displacement of this subset of points is found (arrow in main and cross in inset). A new region of interest is drawn around the mean from the initial iteration (dashed line). The mean displacement from this subset of points (arrow) is shifted to slightly more positive values than the previous mean. At the final iteration, the tabulated mean (arrow) is equivalent to the starting point (dashed line) because the peak is centered within the region of interest.

In order to benchmark this mean shift approach, we evaluated the ability of the algorithm to detect known shifts of simulated datasets of a test cell that is either circular in 2D or cylindrical in 3D, as summarized in Supplementary Figs 1-7. Shifts were estimated by both the mean shift algorithm and by NLLS fitting of a Gaussian to the spatial cross-correlation function of the two datasets. The performance of each algorithm was similar for easy cases with a well-defined peak. For harder cases, the mean shift method outperforms cross-correlation fitting, both by locating the peak with somewhat higher precision, and by more reliably finding the peak over other local maxima (2D: Supplementary Figs 1-2; 3D: Supplementary Figs 3-4). For NLLS fitting, performance can be improved through the use of elegant methods to determine start-points ^10^ (2D: Supplementary Fig 2; 3D: Supplementary Fig 4, Methods), although this is not required when using the mean shift method. The mean shift method is more computationally efficient than cross-correlation based methods, largely because Fast Fourier transforms (FFTs) are not computed in the mean shift approach (Supplementary Fig 5). This improvement in speed is enabled through the use of a particularly efficient algorithm from the R package spatstat ^11^ to extract pairwise distances between nearby points (see Methods).

We have developed a metric that estimates error in the mean-displacement from data, and find that the mean shift algorithm reliably finds the desired peak when the estimated error remains smaller than one quarter of the localization precision (Supplementary Fig 7). The improved performance of the mean shift approach is most apparent in simulated 3D datasets because displacements are identified in all directions simultaneously, rather than using 2D projections with reduced information. Supplementary Fig 6 shows that the full 3D MS algorithm is both more precise and finds the true shift more reliably than the 2D MS algorithm performed on the same data projected into the xy plane. NLLS fitting can also be accomplished in 3D in principle, although computing the cross-correlation in 3D is typically not practical due to the large memory required to tabulate 3D FFTs of images at high spatial resolution. The mean shift approach uses lists of localizations rather than reconstructed images, therefore memory requirements do not increase meaningfully with the dimensionality of the dataset.

This mean shift approach is applied to experimentally obtained SMLM localizations by distributing localizations into temporal bins with equal numbers of frames, and displacement estimates and expected errors are tabulated between all possible pairs of bins. A weighted linear least squares fitting algorithm is then used to generate a trajectory that passes through control points positioned at times centered on each temporal bin, as described previously ^10^.

Drift-corrected reconstructions of Nup210 labeled nuclear pore complex (NPC) assemblies in primary mouse neurons are illustrated in Fig 2a-d. Fig 2b is a reconstruction produced without drift correction in which localizations from single NPCs are smeared over a large area, highlighting the importance of drift correction. The performance of the mean shift algorithm was tested on this dataset by generating multiple drift trajectories through binning with different temporal resolutions. These trajectories were each applied to the full SMLM dataset and the Fourier Ring Correlation (FRC) ^12,13^ was used to quantify image resolution (Fig 2e). The FRC resolution achieves a minimum for temporal bins of <5s, below which the mean shifts regularly fail and the drift correction loses accuracy. This is somewhat below the criterion from simulations, likely because the additional structure in the image assists in finding the global maxima. We conducted similar drift corrections using a NLLS approach, applying two different methods to define the start-point for least-squares fitting. The mean shift algorithm out-performs both cross-correlation approaches, allowing for accurate drift correction with smaller temporal bins, improving the resolution of the reconstructed image. Additional diagnostics are shown in Supplementary Fig 8 for the MS and NLLS approaches.

**Figure 2.**
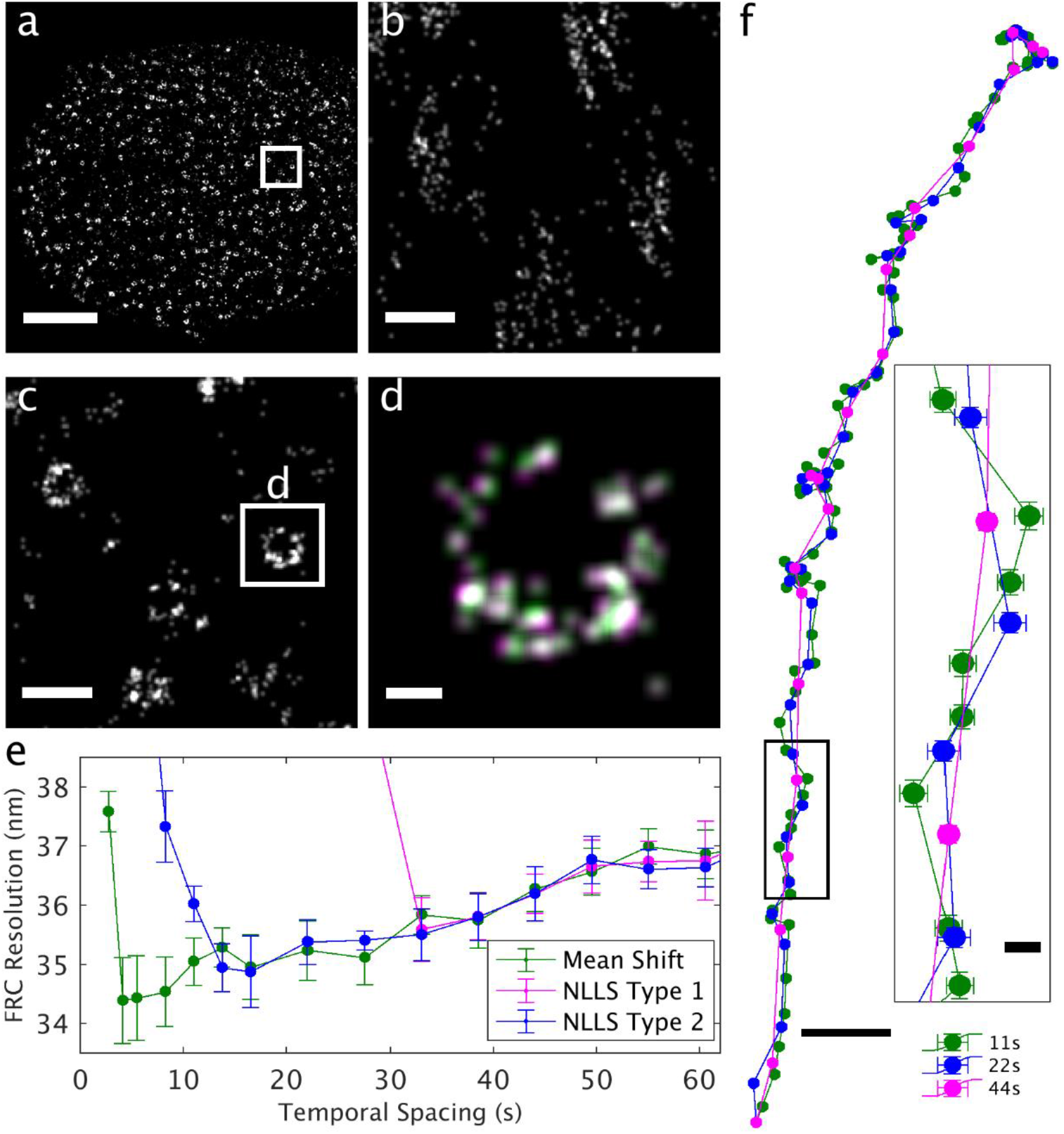
Demonstration of mean shift drift correction of a 2D SMLM dataset of antibody-labelled Nup210 in nuclear pore complexes, within the nuclear envelope of primary mouse neurons. This dataset contained 15500 image frames acquired over 14 minutes, with an average localization precision of 8nm. a) Reconstructed image of a single nucleus which is a subset of this dataset. b) Reconstruction without drift correction of the region shown within the white box in A. c) Same region as in B but with drift correction. d) An image of a single complex demonstrates the modest shifts in localizations due to drift corrections estimated with 11s (green) and 44s (magenta) temporal bins. Scale bars are 2µm (a) 200nm (b,c) and 30nm (d). e) Fourier Ring Correlation (FRC) resolution after applying drift corrections estimated using the specified temporal bin widths for the mean shift and cross-correlation methods. Error bars represent the standard deviation over 20 replicates of the FRC calculation. f) Estimated drift trajectories evaluated from using the method and temporal spacing specified. Error bars represent 68% confidence intervals from the least squares drift estimation of each control point. Scale bars are 20nm for the overall drift curve and 2nm in the inset.

Drift trajectories are shown in Fig 2f for temporal bin-widths that produce accurate FRC metrics for the three approaches. In all cases, a temporal bin slightly larger than the minimum from the FRC curve is used, since this produces smaller errors on individual control points. As expected, the three drift trajectories follow the same general shape but the trajectory generated from the mean shift has improved time resolution. In parts of the trajectory, the errors of the control points are smaller than the distance between the trajectories. In these regions, higher time-resolution yields improved spatial resolution in the final reconstructed image. While the differences in the trajectories are significant, their impact is not apparent when viewing reconstructed images of entire nuclei or collections of NPCs, as in Fig 2a,c. Differences become apparent in images of individual pores, where displacements of several nm shift the relative positions of labeled subunits (Fig 2d).

Figure 3 applies a similar analysis to a SMLM dataset of labeled B cell receptors on the ventral membrane of B cells imaged using a phase mask in the emission path to localize fluorophores in 3D ^14^. Here, the differences in the performance of the mean shift and NLLS fitting methods are more pronounced than in the 2D dataset of Fig 2, as was also seen in simulated data (Supplementary Fig 7). This is largely due to the use of 2D projections to determine displacements by NLLS fitting, while the mean shift approach determines drift in 3 dimensions simultaneously. Additional diagnostics are shown in Supplementary Fig 9.

**Figure 3.**
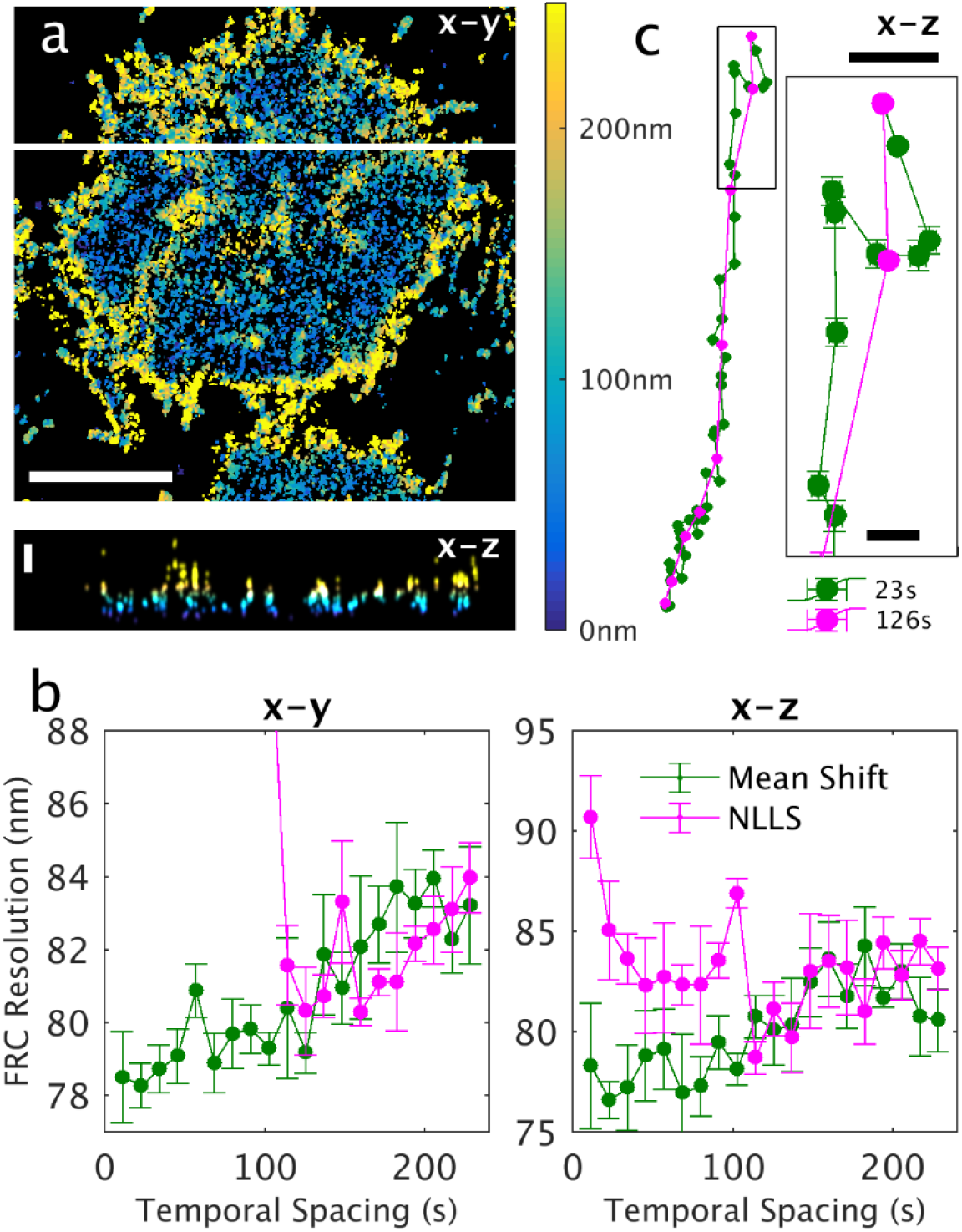
Demonstration of mean shift drift correction of a 3D SMLM dataset of B cell receptors at the ventral plasma membrane of CH27 B cells. This dataset contained more than 400,000 localizations acquired over 15 minutes, with an average localization precision of 17 nm in the lateral (x-y) dimension and 31 nm in the axial (z) dimension. a) Reconstructed image of a subset of this dataset showing the average z position within each x-y pixel as indicated in the color bar. x-z slice at the position drawn as a white line shown below. Scale bars are 5µm for x-y and 200nm for z. b) Fourier Ring Correlation (FRC) estimates of image resolution after applying drift corrections estimated using the specified temporal bin widths for the mean shift and NLLS methods. Error bars represent the standard deviation over 5 replicates of the FRC calculation. (C) Estimated drift trajectories evaluated with the specified temporal spacing. Error bars represent 68% confidence intervals from the least squares drift estimation of each control point. Scale bars are 50nm and 10nm in the inset.

In summary, a mathematically simple mean shift algorithm modestly out-preforms cross-correlation based estimates of drift correction in 2D and more significantly improves the time-resolution of drift-corrections in 3D. The approach is computationally efficient, is robust without sophisticated methods to estimate start-points, and does not require image reconstruction with memory and pixelation limitations. The metric provided to estimate error and predict robustness directly from data provides users with a means to evaluate the quality of a drift correction within an SMLM analysis pipeline. For the example datasets explored, modest improvements in resolution lead to adjustments of localized molecule positions relevant for evaluating the structure of protein complexes in cells.

## Methods

### Simulated Datasets

An idealized 2D SMLM dataset was simulated as a spatially random set of fluorophores on a 20µm diameter circular cell, with each fluorophore giving rise to a Poisson-distributed number of localizations with isotropic Gaussian localization error 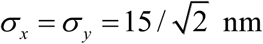. We define 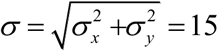 to denote the total root-mean-square localization error. A second dataset was generated from the same fluorophore locations, localization precision, and average number of localizations per fluorophore, and shifted between 0 and 150 nm in a random direction.

Simulated datasets were generated over a range of densities (5 to 100 per µm^2^) and a range of localizations per molecule (.05 to .2).

An idealized 3D SMLM dataset was simulated in a similar fashion. Fluorophores were distributed uniformly on a cylinder 20 µm in diameter and 2 µm deep. Each fluorophore produces a Poisson-distributed number of localizations with 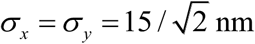 as before, and with 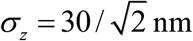. One dataset is translated by a random distance between 0 and 150 nm in a random direction in x, y, and z.

### Extracting Close Pairs of Coordinates Between Datasets

Consider two point sets **u**_*i*_ = (*u*_*ix*_, *u*_*iy*_) and **v** _*j*_ = (*v* _*jx*_, *v* _*jy*_), for *i* = 1,…, *n*_1_ and *j* = 1,…, *n*_2_. We wish to quickly determine which pairs (*i, j*) are closer than some maximum distance *r*_max_ ; i.e. which pairs satisfy ‖ **u**_*i*_ − **v** _*j*_ ‖ < *r*_max_. The algorithm is adapted from the code for the `closepairs()` and `crosspairs()` functions of the R package spatstat ^11^, and implemented in C with a MATLAB interface. We first sort each dataset with respect to its *x*-coordinate, so that *u*_*kx*_ ≤ *u*_*lx*_ whenever *k* ≤ *l*. Then the algorithm proceeds as follows:

1. Let *i* = 1and *j*_left_ = 1.
2. Let *x*_left_ = *u*_*ix*_ − *r*_max_. All close pairs of **u**_*i*_ must satisfy *v* _*jx*_ > *x*_left_.
3. Increment *j*_left_ until 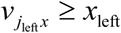.
4. For each *j* = *j*_*left*_, …, *n*_2_, if *v*_*jx*_ − *u*_*ix*_ > *r*_max_, increment *i* and return to step 2. Otherwise, compute 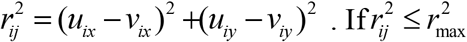. add (*i, j*) to the list of results.

This algorithm avoids computing pairwise distances between most pairs in the dataset, and so is much faster and more memory efficient than a brute force approach. It can be readily adapted to higher dimensions by applying the appropriate *n*-dimensional distance metric in step 4. For convenience, our implementation returns the displacements Δ**r**_*ij*_ = **v** _*j*_ − **u**_*i*_, and total distance *r*_*ij*_ =‖ Δ**r**_*ij*_ ‖ for each pair (*i, j*), instead of the indices themselves.

### Determining Shifts between Translated Datasets Using a Mean Shift Algorithm

Given the set of displacements **Δr**_*ij*_ = **u**_*i*_ − **v** _*j*_ between two point sets **u**_*i*_ and **v** _*j*_, a mean shift clustering algorithm ^5–7^ can be applied to search for the peak of the displacement density function. Briefly, let **r**_shift,0_ be an initial guess to initialize the shift estimate, and δ a radius of consideration to use in the optimization procedure. The algorithm proceeds by iteration, by setting

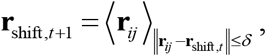

where the average is restricted to the subset of displacements **r**_*ij*_ that satisfy the subscript, i.e. that are within a radius δ from the previous shift estimate **r**_shift,*t*_. The algorithm terminates when the distance ‖**r**_shif*t*, *t* +1_ − **r**_shift,*t*_ ‖ between subsequent shift estimates becomes smaller than machine precision, or when the number of iterations exceeds a user-defined maximum number. δ must be sufficiently large so that the true shift resides within the explored area when centered at the starting-point. In practice, we apply the algorithm twice: first with a large δ to determine the rough shift, and then with a smaller δ, using the first estimate as a starting point, to refine the estimate.

While the above can be applied directly to 3-dimensional data by taking the average over a 3-dimensional ball of radius δ instead of the 2-dimensional disc, we find it is advisable to consider an ellipsoid that is stretched in the z-direction, to account for the larger axial localization errors present in our 3-dimensional simulated and experimental datasets. In the present work, we let the semimajor axis of the ellipsoid be 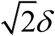, in the z direction, and the semiminor axes both δ, so that x-y cross-sections of the regions of consideration are discs.

### Estimates of mean shift error

We model the distribution of pairs around the true shift as a Gaussian-distributed peak with standard deviation *ζ*, centered on the true shift **r**_shift_ on a uniformly distributed background. Assuming **r**_shift,*t*_ is sufficiently close to **r**_shift_ that most of the Gaussian peak falls within the region of consideration, the variance *ξ* ^2^ of the two components of **r**_shift,*t*_ is given by

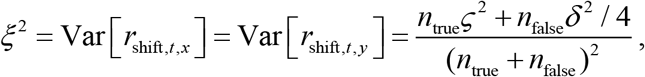

where δ is the radius of consideration for the MS algorithm, and *n*_true_ and *n*_false_ are respectively the number of “true pairs” that are drawn from the Gaussian part of the distribution (displacements between different localizations of the same molecules) and the number of “false pairs” that are drawn from the uniform part (displacements between different molecules), that fall within the region of consideration. Furthermore, the expected value after one more step can be derived:

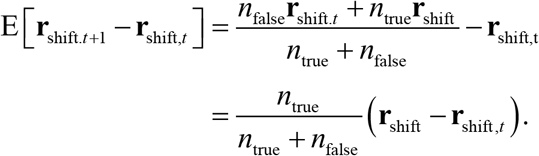

Suppose *t* is the final step of the algorithm, i.e. **r**_shift,*t* +1_ − **r**_shift,*t*_ = 0. Then by hypothesis, **r**_shift,*t*_ deviates from its expected value by

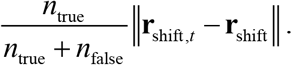

This deviation will typically take on values comparable to the standard deviation *ξ* shown above. Thus, we estimate the error of the MS algorithm by:

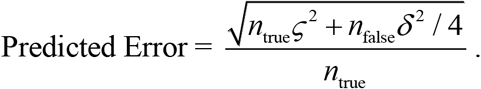

This predicted error is to be interpreted as an estimate of the standard deviation of the shift estimate in each direction.

In practice, the parameters *n*_true_, *n*_false_, and *ζ* are not known, so we estimate them from data. Specifically, we construct the isotropic cross-correlation function *c*(*r*) from the pair separations **r**_*ij*_, determine the baseline of *c*(*r*) from its long-range median value, and use the baseline to infer *n*_true_ and *n*_false_. Finally, we fit *c*(*r*) to a Gaussian plus a constant to estimate *ζ*. This error estimate is derived from a heuristic argument and is not exact. However, its performance is adequate in practice. See Supplementary Fig 7a for a comparison to observed standard deviations of MS shift estimates.

For 3D data, we compute lateral and axial predicted errors separately, by projecting the data from the ellipsoidal region of consideration into the x-y plane or onto the z axis, respectively. *n*_true_, *n*_false_, and *ζ* are estimated separately for the lateral and axial directions from the two projections.

### Evaluating displacements using nonlinear least squares (NLLS) fitting

Displacements **r**_shift_ between pairs of localization datasets were also estimated by NLLS fitting of a Gaussian to the spatial cross-correlation function of the two datasets. NLLS fitting was either accomplished using custom software written in Matlab used previously by our group or with software published as supplementary material of ^10^. For both NLLS approaches, images were first reconstructed from simulated localizations with a pixel size of 15nm for simulated localizations, or from acquired data with a pixel size of 8nm for Nup210 or 15nm for B cell receptor experimental localizations. Both codes then tabulate cross-correlations using 2D Fast Fourier Transforms then fit a 2D Gaussian function to a subset of the cross-correlation centered at the start-point of the NLLS algorithm. In our software, the starting point for the least squares fitting was determined by fitting the whole dataset to a broad Gaussian shape. The software from ^10^ finds the start-point using an elegant smoothing step to reduce noise then uses the largest local maximum of the smoothed cross-correlation as the start-point for fitting. The choice of start-point is important in NLLS to prevent the algorithm halting at a local maximum. The ‘failures’ of Supplementary Figs 1-4 are in most cases examples of such local optima. In all cases, the smoothing, peak-finding algorithm of ^10^ outperforms our pre-fitting approach.

For localizations acquired in 3D multiple 2D projections were constructed from 3D localizations, then the procedures described for 2D images were applied to determine displacements. First, images projecting on the lateral dimension (x-y plane) were generated and the lateral displacement was determined. To compute the z displacement, both the xz and yz projections were used, and the final z displacement was the average determined from the two projections. Computation time was assessed in MATLAB using the built-in `tic` and `toc` functions. For display in Supplementary Fig 5, computation time was averaged over 500 simulations for each condition.

### Correcting continuous drift

Continuous drift was corrected by temporally dividing the data into *N* bins, each having the same number of frames. For each of the *N* (*N* −1) / 2 pairs (*m, n*) of temporal bins, the mean shift or NLLS algorithm is applied to estimate the shift **r**_shift,*m*→*n*_ from temporal bin *m* to *n*, corresponding to the drift between the bins. Drift at each of the *N*time points is calculated from the *N*(*N*− 1)/2 pairwise shifts using a least-squares minimization algorithm ^10^; this takes advantage of the overdetermined nature of the drift calculation to improve the precision of the measurement. Outlier shifts whose residual with respect to the least-squares estimate exceeds a user-defined threshold, can also be discarded as described in ^10^. These shifts typically correspond to “failures” of the shift estimation method. The final drift curve at each frame is determined by linear interpolation and extrapolation from the *N* basis points.

### Evaluating the resolution of drift-corrected datasets

Resolutions of the final reconstructed images were compared using Fourier Ring Correlation ^12,13^. To compute the x-y resolution of the nuclear pore complex dataset, nearby localizations belonging to adjacent camera frames were grouped together, with the position taken to be the average of the relevant coordinates. The FRC curves were produced by dividing the dataset into blocks of 500 frames and allocating an equal number of blocks randomly to each of the two sets. The pixel size was taken to be 2.5 nm. To compute the resolution of the B cell dataset, the Fourier Ring approach was applied to the xy and xz projections in turn, using a pixel size of 5 nm in each case.

### Preparation of cellular samples for imaging

Mouse primary neurons were isolated from P0 mouse pups that were decapitated and brains were isolated into ice cold, filtered dissection buffer (6.85 mM sodium chloride, 0.27mM potassium chloride, 0.0085mM sodium phosphate dibasic anhydrous, 0.011mM potassium phosphate monobasic anhydrous, 33.3mM D-glucose, 43.8mM sucrose, 0.277mM HEPES, pH 7.4) as described in ^15^. After removing the cerebellum and the meninges, cortices were dissected out, placed into a microcentrifuge tube, and cut into small pieces with dissection forceps. Cortices were incubated in 50µL papain (2mg/mL; BrainBits) and 10µL DNase I (1mg/mL; Worthington Biochemical) for 30min at 37 °C. 500µL BrainPhys Neuronal Medium (Stemcell Technologies) and 10µL additional DNase I were added, and cortices were titurated using P1000 and P200 pipet tips. Titurated cortices were centrifuged at 1000rpm for 5min. After discarding the supernatants, the pellets were titurated and centrifuged three more times until the supernatant remained clear and neuronal pellets were visible. Pelleted neurons were resuspended in BrainPhys Neuronal medium with SM1 supplement as previously described ^16^, then plated onto 35mm #1.5 glass-bottom dishes (MatTek Life Sciences) coated with polyethlenimine (100 µg/ml; Polysciences). Neurons were incubated in 5% CO_2_ at 37 °C, and 1mL of media was replaced every four days.

On day 10 of culture (*days in vitro* 10), neurons were rinsed with sterile Hank’s Balanced Salt Solution, then incubated for 1min with pre-warmed 2% PFA (Electron Microscopy Sciences) in Phosphate Buffered Saline (PBS). The neurons were then incubated in 0.4% Triton X-100 (Millipore Sigma) in PBS for 3min, and fixed for 30min with 2% PFA in PBS. Neurons were then washed with PBS five times, incubated in blocking buffer containing 5% Normal Donkey Serum and 5% Normal Goat Serum (Jackson Laboratories) for 30min, then labeled with Nup210 polyclonal antibody diluted in blocking buffer (1:200; Bethyl laboratories A301-795A) overnight in 4 °C. The following day, neurons were washed three times in PBS then stained with Goat-anti-rabbit Alexafluor 647 secondary antibody (1:1000; Thermo Fisher) for an hour, washed three times with PBS, then imaged.

CH27 B cells ^17^ were cultured, allowed to adhere to 35mm #1.5 glass-bottom dishes (MatTek Life Sciences) overnight, then incubated in Alexa647 conjugated fAb prior to fixation in 4% PFA and 0.1% gluteraldehide (Electron Microscopy Sciences), as described previously ^18^. The labeled fAb antibody was prepared by conjugating an Alexa647 NHS ester (ThermoFisher) to an unconjugated fAb (Goat Anti-Mouse IgM, µ chain specific; Jackson Immunoresearch) using established protocols ^18^.

### Single molecule image and localization

Imaging was performed using an Olympus IX83-XDC inverted microscope. TIRF laser angles where achieved using a 60X UAPO TIRF objective (NA = 1.49), and active Z-drift correction (ZDC) (Olympus America) as described previously. The ZDC was not used for collection of 3D datasets. Alexa 647 was excited using a 647 nm solid state laser (OBIS, 150 mW, Coherent) coupled in free-space through the back aperture of the microscope. Fluorescence emission was detected on an EMCCD camera (Ultra 897, Andor) after passing through a 2x expander. Imaging in 3D was accomplished using a SPINDLE module equipped with a DH-1 phase mask (DoubleHelix LLC).

Single molecule positions were localized in individual image frames using custom software written in Matlab. Peaks were segmented using a standard wavelet algorithm ^19^ and segmented peaks were then fit on GPUs using previously described algorithms for 2D ^20^ or 3D localizations ^21^. After localization, points were culled to remove outliers prior to drift correction. Images are rendered by generating 2D histograms from localizations followed by convolution with a Gaussian for display purposes.

## Acknowledgements

This work was supported by grants from the U.S. National Science Foundation (MCB1552439) and National Institutes of Health (R01GM129347).

## Author Contributions

The method was devised by FJF and TRS in consultation with SLV. FJF wrote the majority of the code, and performed the majority of analyses in consultation with TRS and SLV. FJF, TRS and SLV wrote the text. SK prepared and imaged nuclear pore complex samples for Fig 2. RAB prepared and imaged B cell samples for Fig 3.

## Supplementary Software

Supplementary Software can be found at https://github.com/VeatchLab/Mean-Shift-Drift-Correction. The software contains Matlab and C code to run mean shift drift corrections on 2D and 3D SMLM data. Three example scripts are also included:

1. meanshift_example.m: determine the shift between a single pair of sets of localizations sampling the same structure at different times. The example uses data from the 2D nuclear pore complex (NPC) dataset of Fig 2.
2. example_NPC.m: correct 2D drift from the full NPC dataset of Fig 2.
3. example_Bcell.m: correct 3D drift from the full B cell dataset of Fig 3.

## Supplementary Figures

**Supplementary Figure 1.**
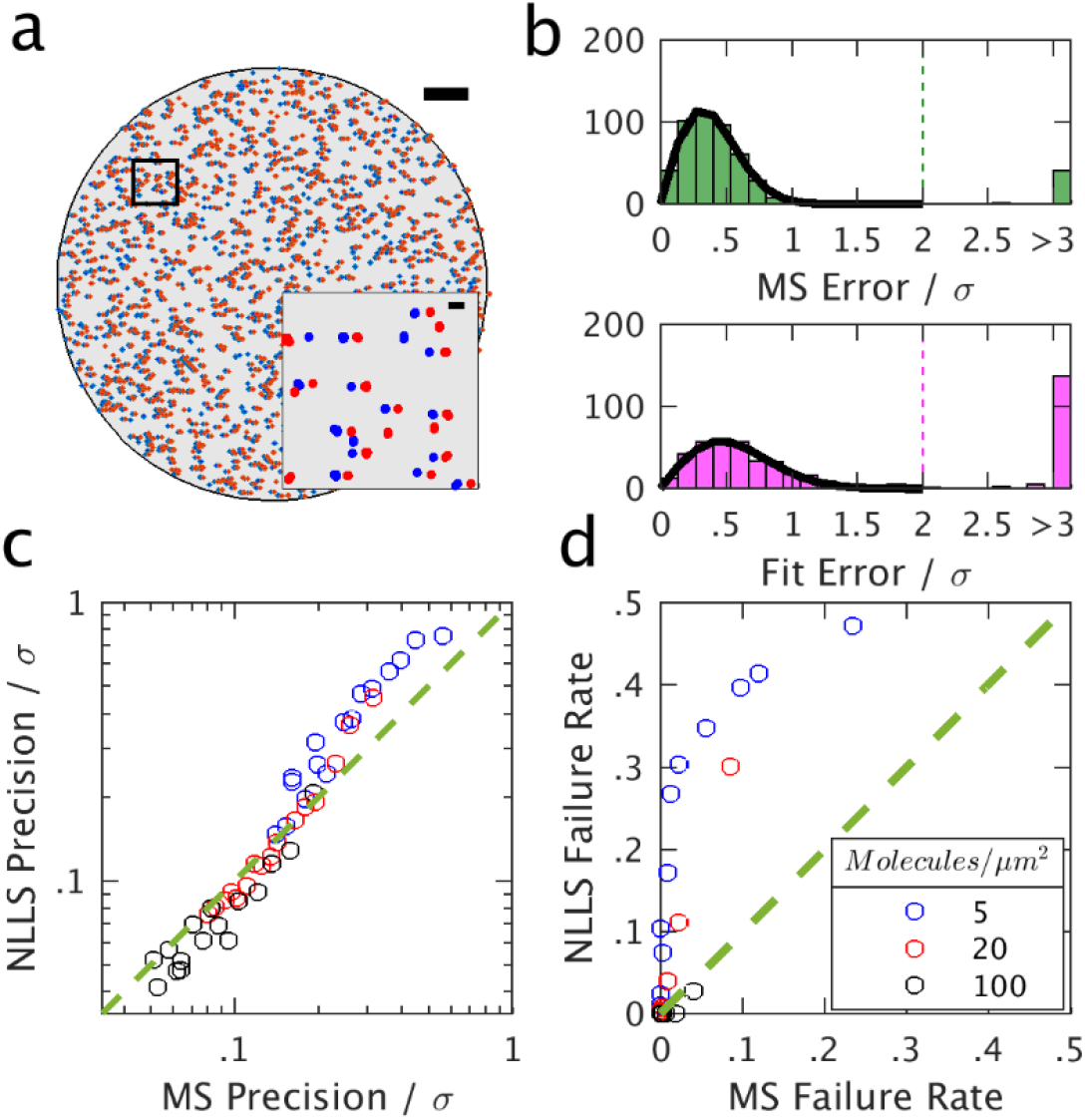
Evaluating the mean shift (MS) algorithm on 2D simulated data, compared to the NLLS “Type 1” approach. Simulations and shift determination approaches are described in methods. **a)** Example simulated SMLM dataset. Scale bars are 2 µm (large image) and 150 nm (inset). **b)** Histograms of errors for the MS and NLLS approaches for the 20 molecules/µm^2^, 0.05 localizations per molecule condition. The precision of each method is evaluated for each condition as the standard deviation of a Gaussian fit to the central peak of the histogram. The “failure rate” is the fraction of simulations whose error exceeds twice the localization precision σ, indicated as a dashed line. **c**,**d)** Comparison of the performance of the MS and NLLS approaches. Each point corresponds to 500 simulations of a particular density/overcounting scenario. The green dashed lines indicate equal performance. **c)** Comparison of the standard deviation of the error of each approach, determined by fitting as in (b). **d)** Comparison of failure rates of each approach, as determined in (b).

**Supplementary Figure 2.**
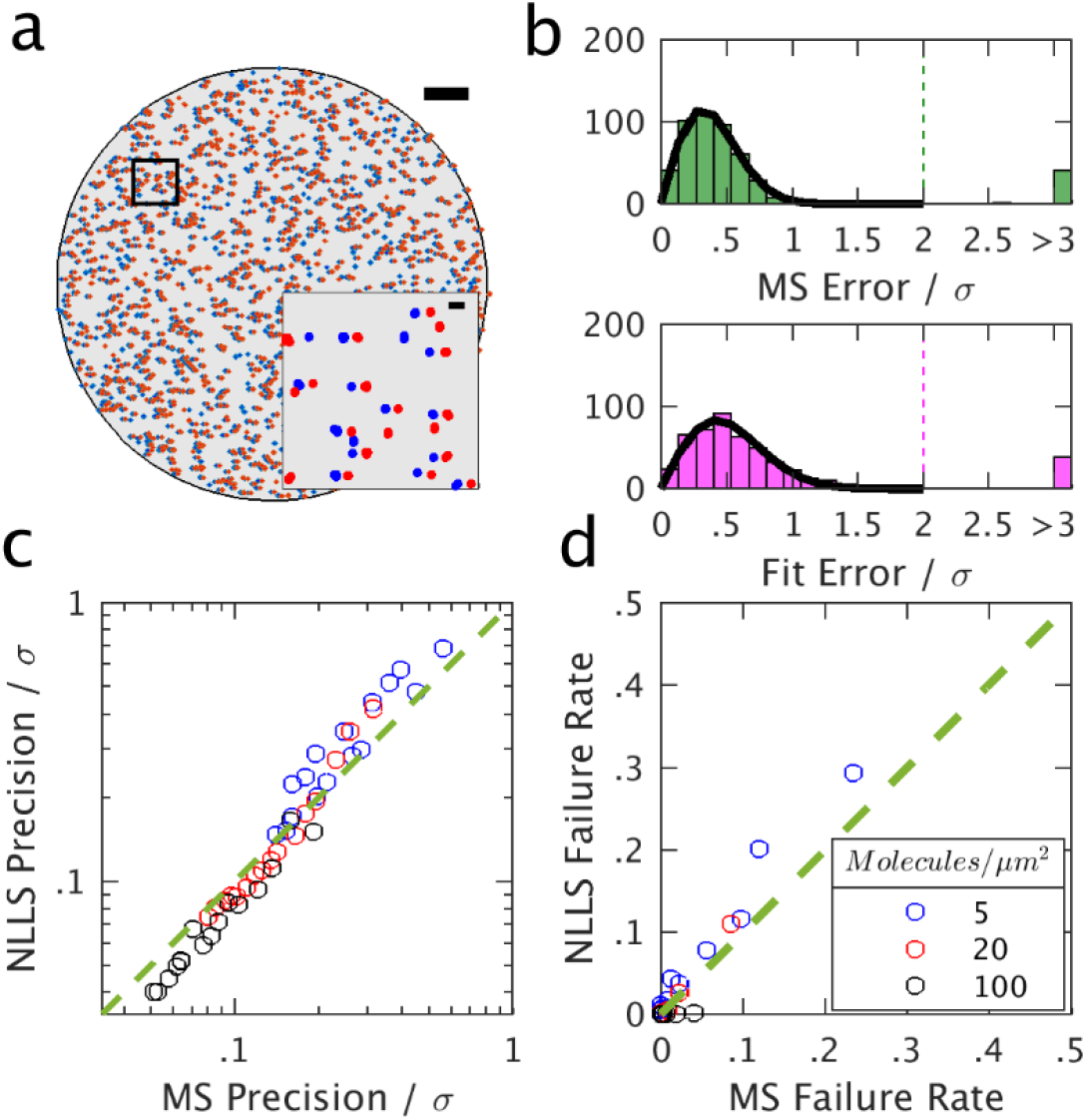
Evaluating the mean shift (MS) algorithm on 2D simulated data, compared to the NLLS “Type 2” approach. Simulations are described in methods. **a)** Example simulated SMLM dataset. Scale bars are 2 µm (large image) and 150 nm (inset). **b)** Histograms of errors for the MS and NLLS approaches for the 20 molecules/µm^2^, 0.05 localizations per molecule condition. The precision of each method is evaluated for each condition as the standard deviation of a Gaussian fit to the central peak of the histogram. The “failure rate” is the fraction of simulations whose error exceeds twice the localization precision σ, indicated as a dashed line. **c,d)** Comparison of the performance of the MS and NLLS approaches. Each point corresponds to 500 simulations of a particular density/overcounting scenario. The green dashed lines indicate equal performance. **c)** Comparison of the standard deviation of the error of each approach, determined by fitting as in (b). **d)** Comparison of failure rates of each approach, as determined in (b).

**Supplementary Figure 3.**
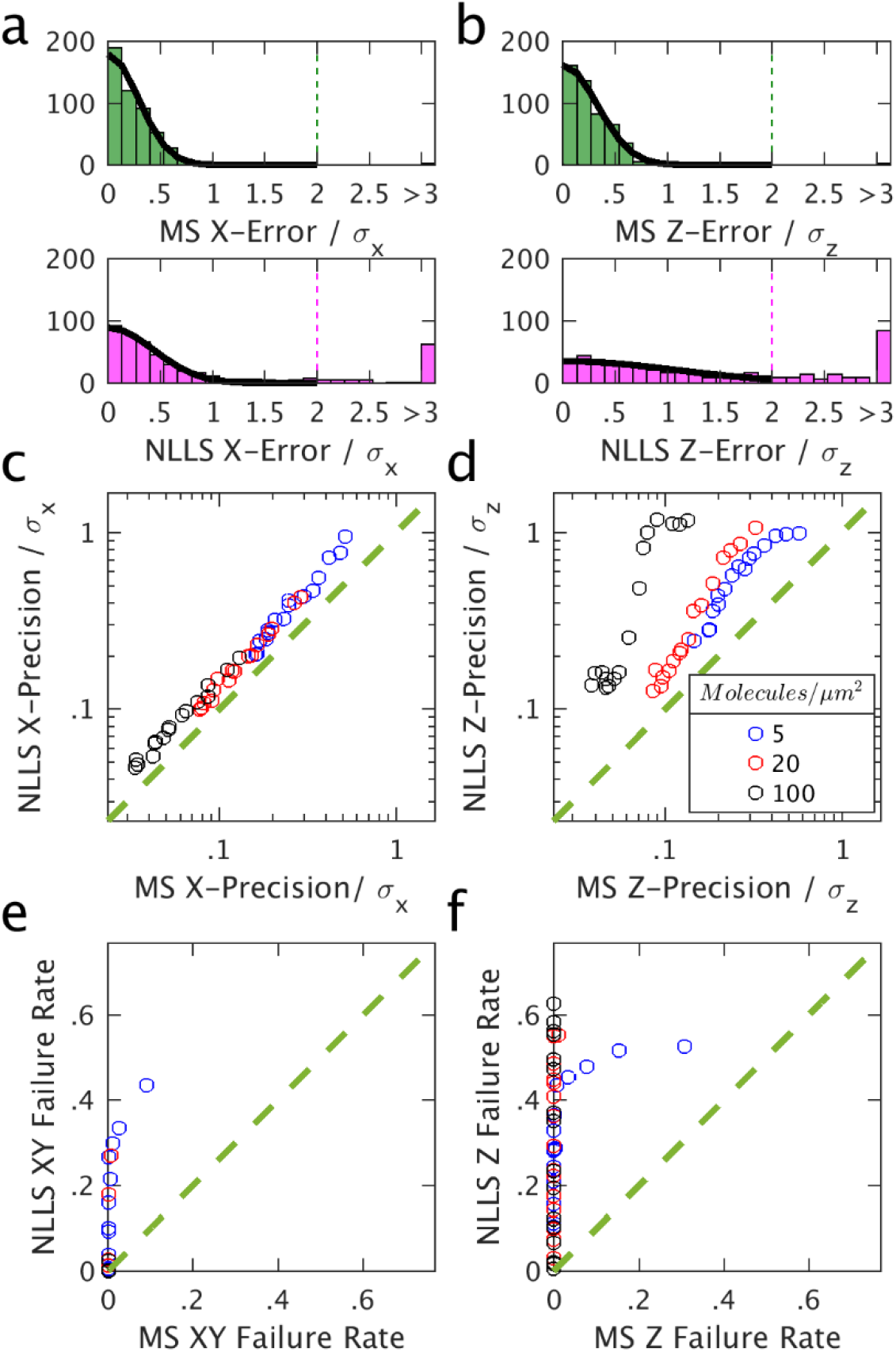
Evaluating the mean shift (MS) algorithm on 3D simulated data, compared to the NLLS “Type 1” approach. See Methods for simulation details. **a,b)** Histograms of x-errors **(a)** and z-errors **(b)** for the MS and NLLS approaches for the 0.01 molecules/µm^3 and 0.05 localizations per molecule condition. The precision of each method is evaluated for each condition as the standard deviation of a Gaussian fit to the central peak of the histogram. The “failure rate” is the fraction of simulations whose error exceeds twice the localization precision σ, indicated as a dashed line. **c,d)** Comparison of the lateral **(c)** and axial **(d)** precision of each approach. The green dashed line indicates equal precision. **e,f)** Comparison of the failure rate of each approach in lateral **(e)** and axial **(f)** directions under the same range of conditions.

**Supplementary Figure 4.**
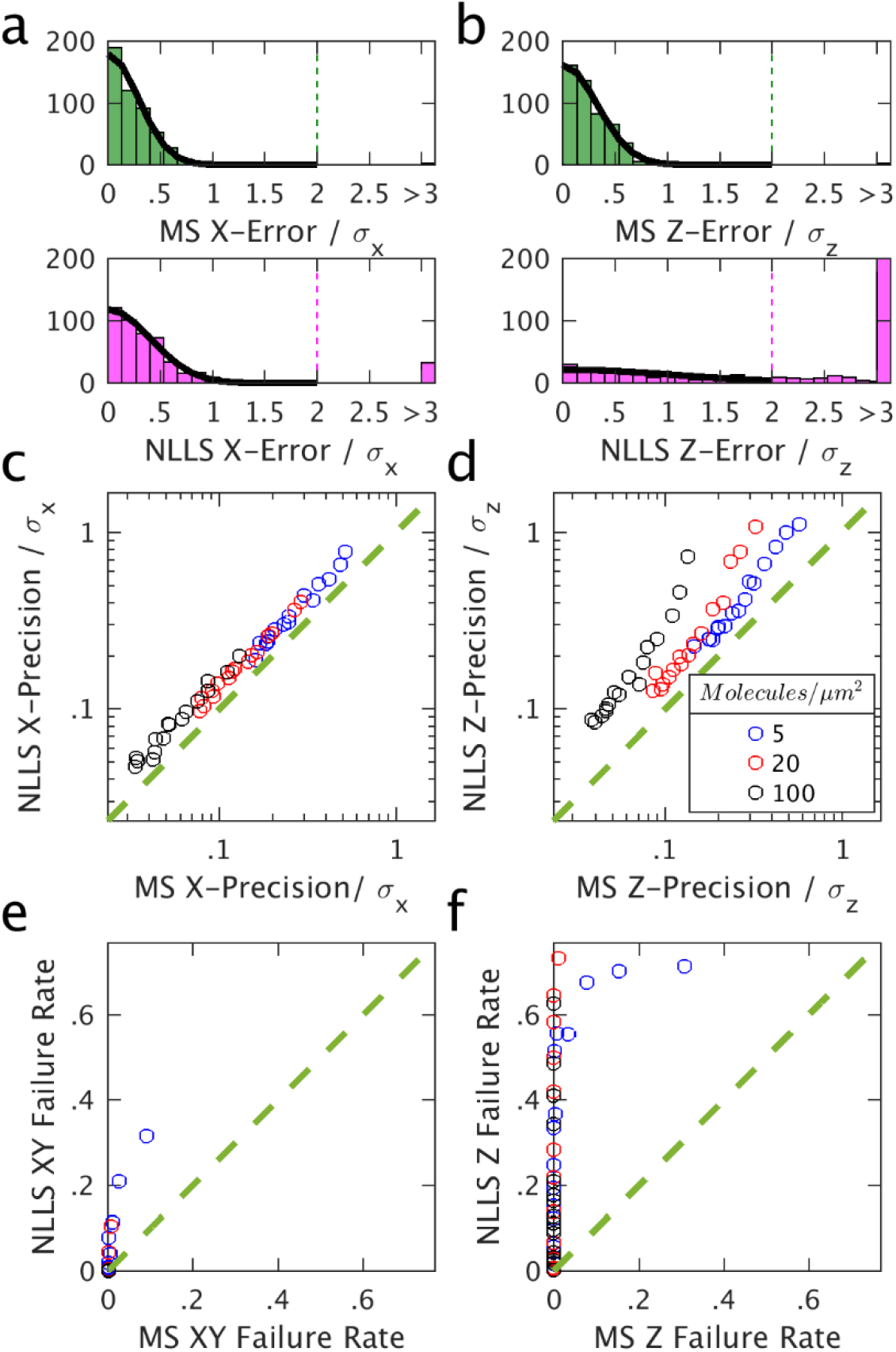
Evaluating the mean shift (MS) algorithm on 3D simulated data, compared to the NLLS “Type 2” approach. See Methods for simulation details. **a,b)** Histograms of x-errors **(a)** and z-errors **(b)** for the MS and NLLS approaches for the 0.01 molecules/µm^3 and 0.05 localizations per molecule condition. The precision of each method is evaluated for each condition as the standard deviation of a Gaussian fit to the central peak of the histogram. The “failure rate” is the fraction of simulations whose error exceeds twice the localization precision σ, indicated as a dashed line. **c,d)** Comparison of the lateral **(c)** and axial **(d)** precision of each approach. The green dashed line indicates equal precision. **e,f)** Comparison of the failure rate of each approach in lateral **(e)** and axial **(f)** directions under the same range of conditions.

**Supplementary Figure 5.**
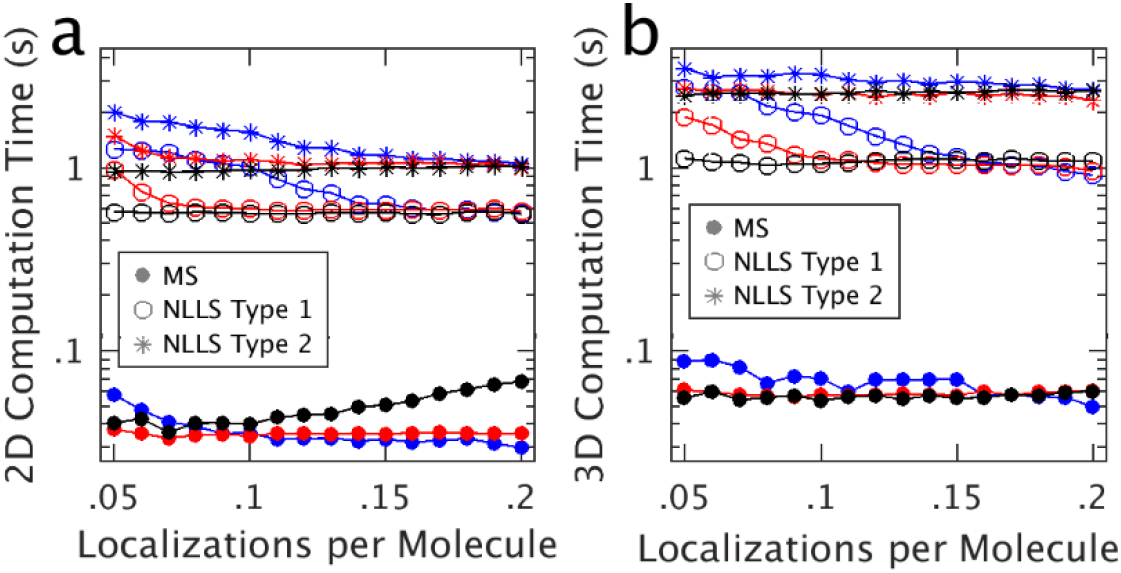
Computation times for mean shift (MS) and nonlinear least squares fitting (NLLS) using two methods to determine starting points for fitting (Type 1 and Type 2) corresponding to the NLLS fitting of Figs S1,3 and S2,4 respectively. **a)** on 2D simulated datasets as in Figs S1,2. **b)** on 3D simulated datasets as in Figs S3,4.

**Supplementary Figure 6.**
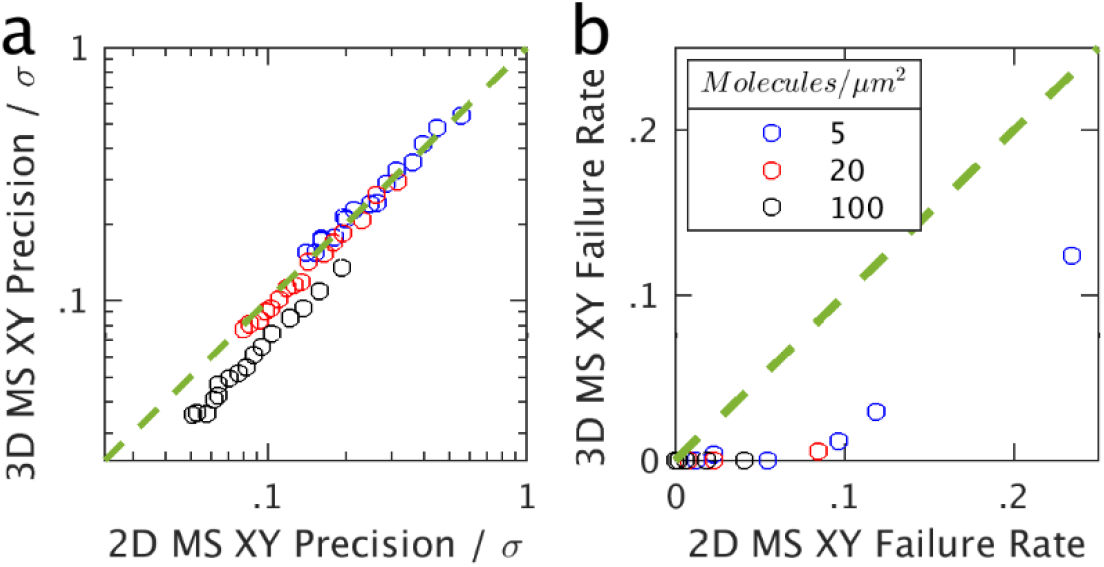
2D projections of 3D data degrade mean shift (MS) shift estimation performance. Lateral (x-y) precision **(a)** and failure rate **(b)** when MS shift is determined in 3D or in 2D after projecting the localizations into the x-y plane.

**Supplementary Figure 7.**
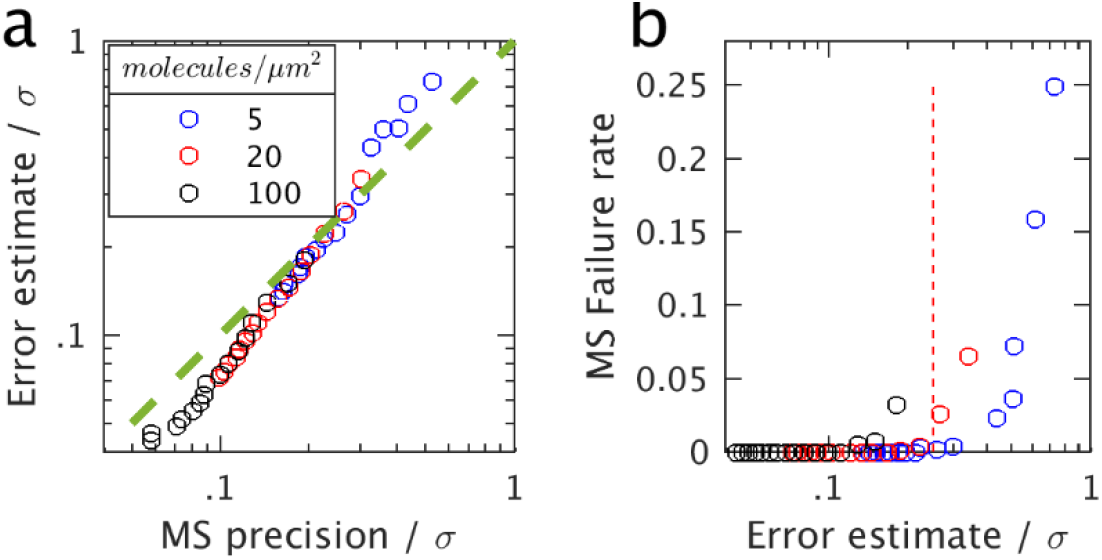
Predicted errors of mean shift (MS) shift estimation. **a)** Comparison of error estimates to observed precision of MS drift estimates, for each simulation condition of Figure S1. **b)** MS failure rate vs error estimate, with the σ/4 criterion shown.

**Supplementary Figure 8.**
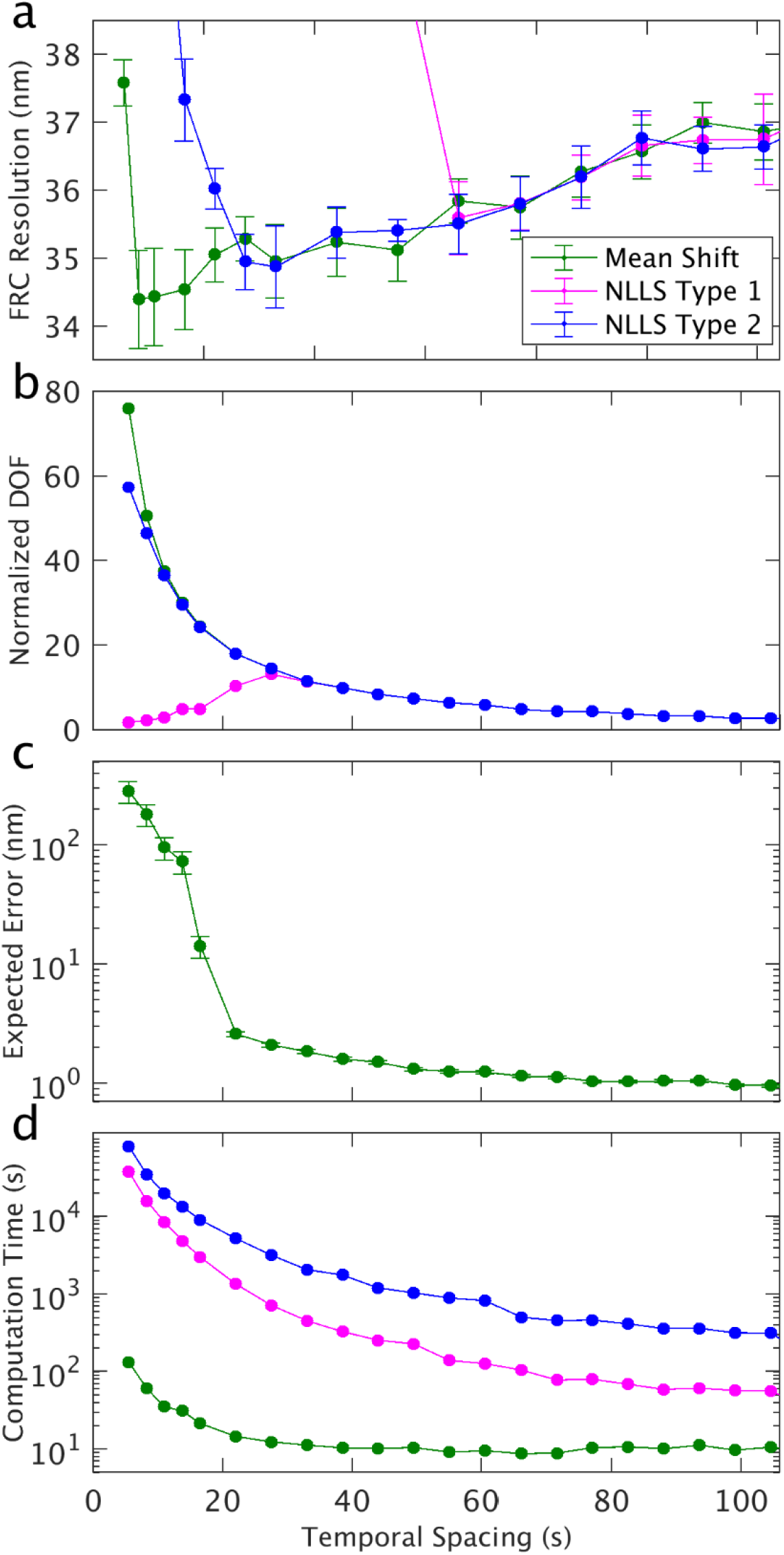
Drift correction diagnostics for the nuclear pore complex dataset of Figure 2. **a)** FRC resolutions. Error bars are given by the standard deviation over 20 trials. **b)** The number of degrees of freedom after removing outliers (normalized by the number of parameters) for the redundant least square minimization calculation. **c)** RMSE of the expected errors for the mean shift method. Error bars are given by the standard error of the mean. **d)** Total computation time.

**Supplementary Figure 9.**
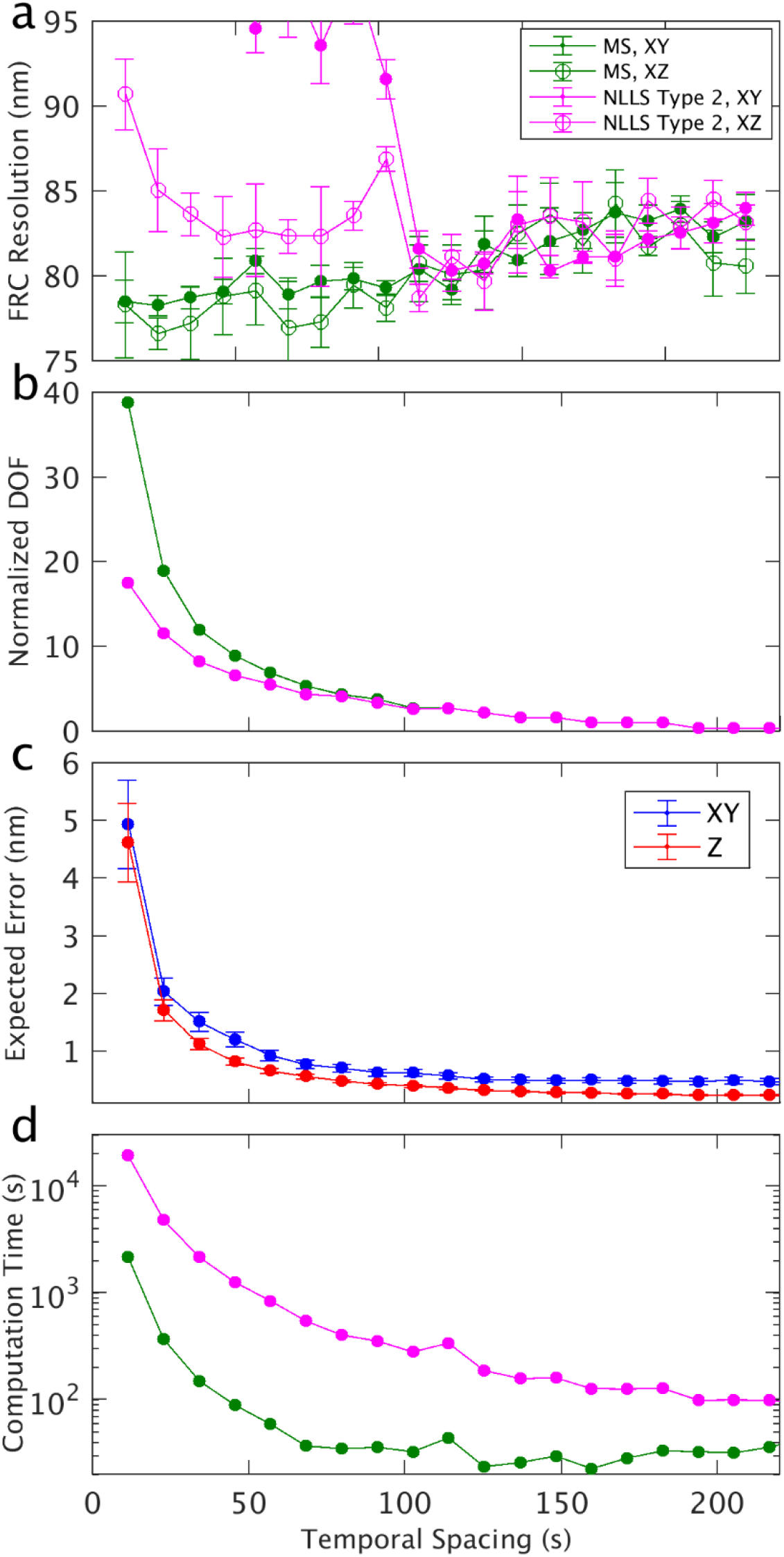
Drift correction diagnostics for the 3D B cell dataset of Figure 3. **a)** FRC resolutions. Error bars are given by the standard deviation over five trials. **b)** The number of degrees of freedom after removing outliers (normalized by the number of parameters) for the least square minimization calculation. **c)** RMSE of the lateral (x-y) and axial (z) expected errors for the mean shift calculation of pairwise shifts. Error bars are given by the standard error of the mean. **d)** Total computation time.

